# Parasite defensive limb movements enhance signal attraction in male little torrent frogs: insight into the evolution of multimodal signals

**DOI:** 10.1101/2021.12.31.474622

**Authors:** Longhui Zhao, Jichao Wang, Haodi Zhang, Tongliang Wang, Yue Yang, Yezhong Tang, Wouter Halfwerk, Jianguo Cui

**Author notes:** Corresponding to Jianguo Cui.

## Abstract

Many animals rely on complex sexual signals that target multiple senses to attract mates and repel rivals. These multimodal mating displays can however also attract unintended receivers, which can be an important driver of signal complexity. Despite being taxonomically widespread, we often lack insight into how multimodal signals evolve from unimodal signals and in particular what roles unintended eavesdroppers play. Here we assess whether the physical movements of parasite defense behavior increase the complexity and attractiveness of an acoustic sexual signal in the little torrent frog (*Amolops torrentis*). Calling males of this species often display limb movements in order to defend against blood-sucking parasites such as frog-biting midges that eavesdrop on their acoustic signal. Through mate choice tests we show that some of these midge-evoked movements influence female preference for acoustic signals. Our data suggest that midge-induced movements may be incorporated into a sexual display, targeting both hearing and vision in the intended receiver. Females may play an important role in incorporating these multiple components because they prefer signals which combine multiple programs. Our results thus help to understand the relationship between ecological and sexual selection pressure operating on signalers and how in turn this may influence multimodal signal evolution.

## Introduction

Sexual selection can drive signal evolution through preferences for complex mating displays ^1-3^. Sexual signalers can either increase their attractiveness by enhancing the complexity in a single sensory modality or by evolving displays that target multiple sensory modalities ^4, 5^. Such multimodal signaling can be highly complex, often involving multiple underlying neuronal motor programs that need to be synchronized in order to perform well ^5-8^. The production and reception of multimodal signals is often more costly in terms of energy loss or increased of predation and parasitism when compared to unimodal signals ^9^, and their evolution is therefore often explained through functional benefits, such as cross-modal perception by receivers, which can improve signal detection and discrimination, or enhance attention and memory time ^7, 10, 11^. We know however far less how multimodal signals evolve from unimodal ones. An important question remains whether and when multimodal signals evolve *de novo*, or evolve through a process of co-option, by incorporating additional cues into a unimodal mating display ^7^.

Most species generate by-product cues during signaling. For instance, floating frogs produce water ripples when calling from the water. These ripple cues have become part of the sexual display, as their presence modulates receiver responses to their acoustic signal components ^12^. Multimodal signals can thus originate from cues associated with primary signal production, either through a physical linkage (e.g. case of call-induced water ripples) or through cues generated by other non-communicative behaviors that have subsequently been integrated as part of a sexual display. Such process of co-option has been proposed for many ritualized visual displays which are predicted to have evolved from different intra- or interspecific activities such as intention movements, protective and autonomic responses ^13, 14^. For example, comparative analyses on Anatidae (i.e. ducks) suggest that the precopulatory displays of head-dipping seem to be derived from bathing behavior ^15^.

Anurans (i.e. frogs) provide a good opportunity to test whether non-communicative behaviors can be co-opted into a sexual display function ^16, 17^. In some anurans, for example, arm-waving movements appear to originate from cleaning behavior due to the similarities in both displays ^18^. Furthermore, many species display defensive movements that are also similar with communicative visual displays ^16, 17^. Here we hypothesized that some of these physical movements originated from anti-parasite behavior in frogs. Amphibians are often confronted with parasitism from a range of different insects, such as mosquitos or midges ^19-23^. These parasites are often attracted to the frog’s mating call to collect a blood meal. In return they may transmit blood-borne diseases or endo-parasites, thus imposing a large cost to a calling frog ^24-26^. Some anurans are observed to perform defensive physical movements in response to these parasites.

Limb movements evoked by host-parasite interactions may increase the complexity of sexual displays and can potentially increase female preference for acoustic signals in frogs. Insect-evoked movements are often similar with limb movements that act as visual sexual signal. Thus, the parasite-host interaction may provide an important source of evolutionary raw material for ritualized visual displays in anurans, which in turn may have led to the evolution of multimodal sexual displays in which visual movements and acoustic calls have been combined.

In the present study we examined the role of insect parasites in driving multimodal signal evolution in the little torrent frog (*Amolops torrentis*), a tropical species that breeds in noisy mountain streams and displays both day and night. Male little torrent frogs prefer to emit advertisement calls from the rocks near streams or near vegetation. Calling males are also often observed to be disturbed by various insects including midges and some other potential parasites. In order to repel these insects, they usually generate limb movements that are similar to some spontaneous movements as well as to visual displays that have been reported in other torrent frogs. These observations suggest that their defensive movements may act as a visual cue component in addition to the acoustic component of the little torrent frog’s mating display.

Here we evaluated whether the presence of eavesdropping parasites increase limb movements and determined whether and how these parasite-evoked physical displays influence female preference for multimodal signals. First, we filmed little torrent frogs in the field and classified their physical displays involving limb movements. Second, we assessed the link between parasitism and male limb display by quantifying the frequency of parasite interactions as well as limb movements, and tested whether calling males produced more parasite-evoked displays compared to silent males. Third, we determined the effect of midge-evoked visual movements on female mate-choice with and without the advertisement calls presented. Finally, we tested whether exaggerated movements play a role in mate choice by analyzing the change of male foot-flagging and leg-stretching displays when females approached them in the field.

## Results

### The diverse repertoire of limb displays

Male little torrent frogs possess a rich repertoire of visual displays involving the movements of limbs. Their definitions and detailed descriptions (modified from previous reports ^14, 27^) were as follows: (1) Toe trembling (TT): Vibrating, wiggling or twitching the toes, with the arm and leg motionless (figure S1A); (2) Hind foot lifting (HFL): Raising one hind foot towards the dorsal direction and then returning it back on the ground, without extending the leg (figure S1B); (3) Arm waving (AW): Lifting one of two arms and waving it up and down in an arc towards the front of head (figure S1C); (4) Limb shaking (LSA): Rapid movements of hand or foot in an up-and-down pattern (figure S1D); (5) Wiping (W): Moving a hand or foot on the ground, with the limb not fully extended (figure S1E); (6) Leg stretching (LS): Stretching one leg or both legs at the substrate level (figure S1F); (7) Foot flagging (FF): Raising one or both legs off the substrate level, extending it/them out and back in an arc shape, and then getting it/them back to the ground (figure S1G). Both LS and FF were movements of the hind limb and they were occasionally performed in a similar pattern that was not easy to distinguish. We therefore categorized this two movements collectively LS + FF throughout the remaining part of this study.

### Parasites induce more limb movements

Five observed limb movements are not only spontaneously generated, but also induced by insects (figure 2*A*). The passive visual movements were predominantly evoked by some potential hematophagous parasites such as midges and sandflies (movie S1, movie S2 and figure S2). Specifically, we identified *Corethrella* spp midges and *Phlebotomus* flies, which prefer to feed on the blood of ectotherms ^24, 28^. As seen in figure 2*A*, the movements that were produced by parasite interactions had a high proportion of visual cues, such as wiping (W, 49.3%), arm waving (AW, 38.8%), limb-shaking (LSA, 32.9%) and hind-foot lifting (HFL, 20.4%), while the proportion was low in the toe-trembling (TT, 12.5%) category. Leg-stretching and foot-flagging were not induced by parasite interactions (LS + FF, 0%; table S1). Interestingly, we observed a positive correlation between the level of parasite interference and the total number of visual movements (Pearson’s correlation; *N* = 39; *R* = 0.830; *P* < 0.001; figure 2*B*). We also ran the same analyses but restricted to the two movements found to be attractive to females (i.e., AW and HFL movements; see also below). As a result, we found the same correlation between AW/HFL and the presence of parasites (Pearson’s correlation; *N* = 39; *R* = 0.737; *P* < 0.001).

**Figure 1.**
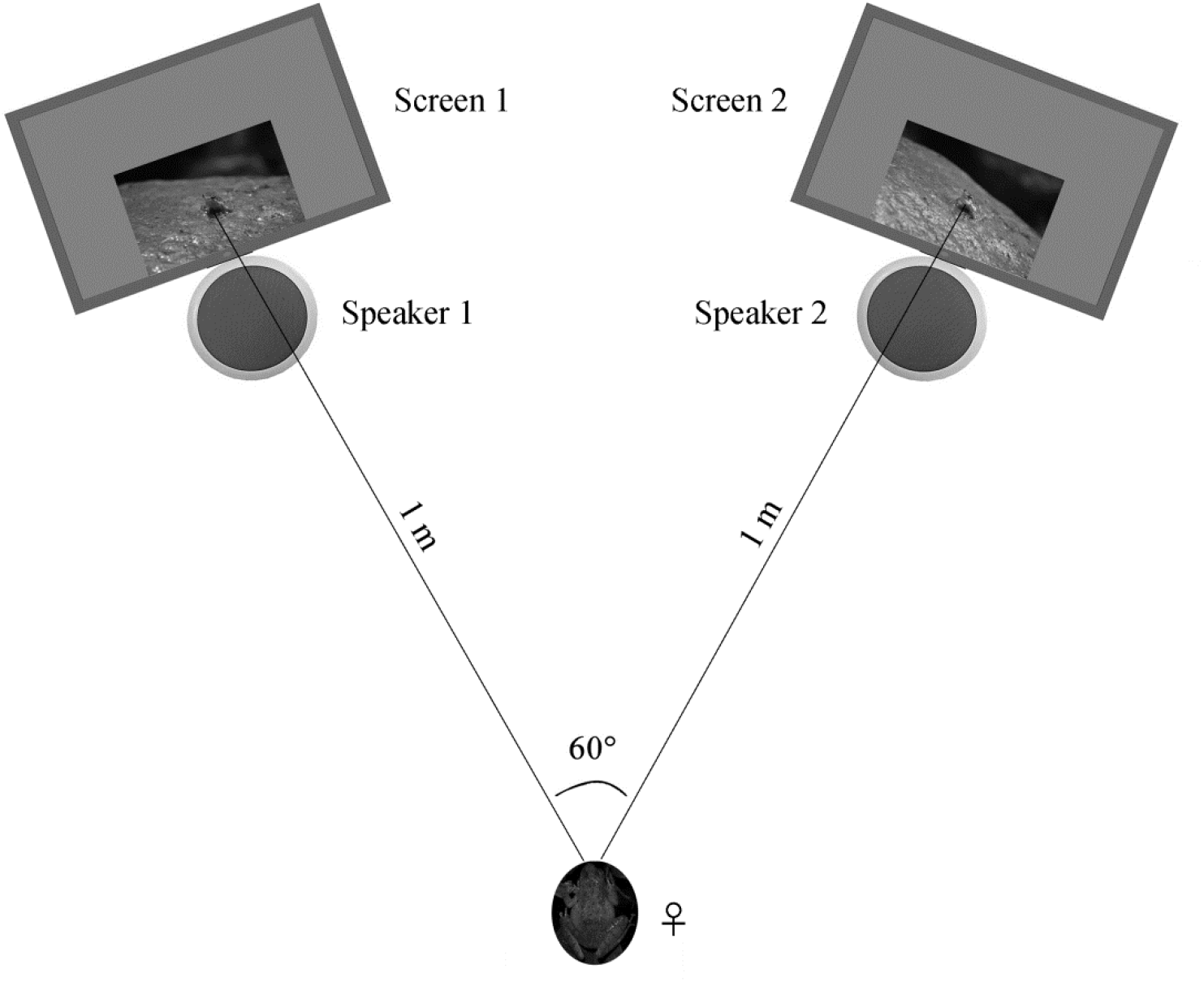
Schematic of the acoustic and visual playback arena. The picture of female frog represents the initial placement point for each playback test.

**Figure 2.**
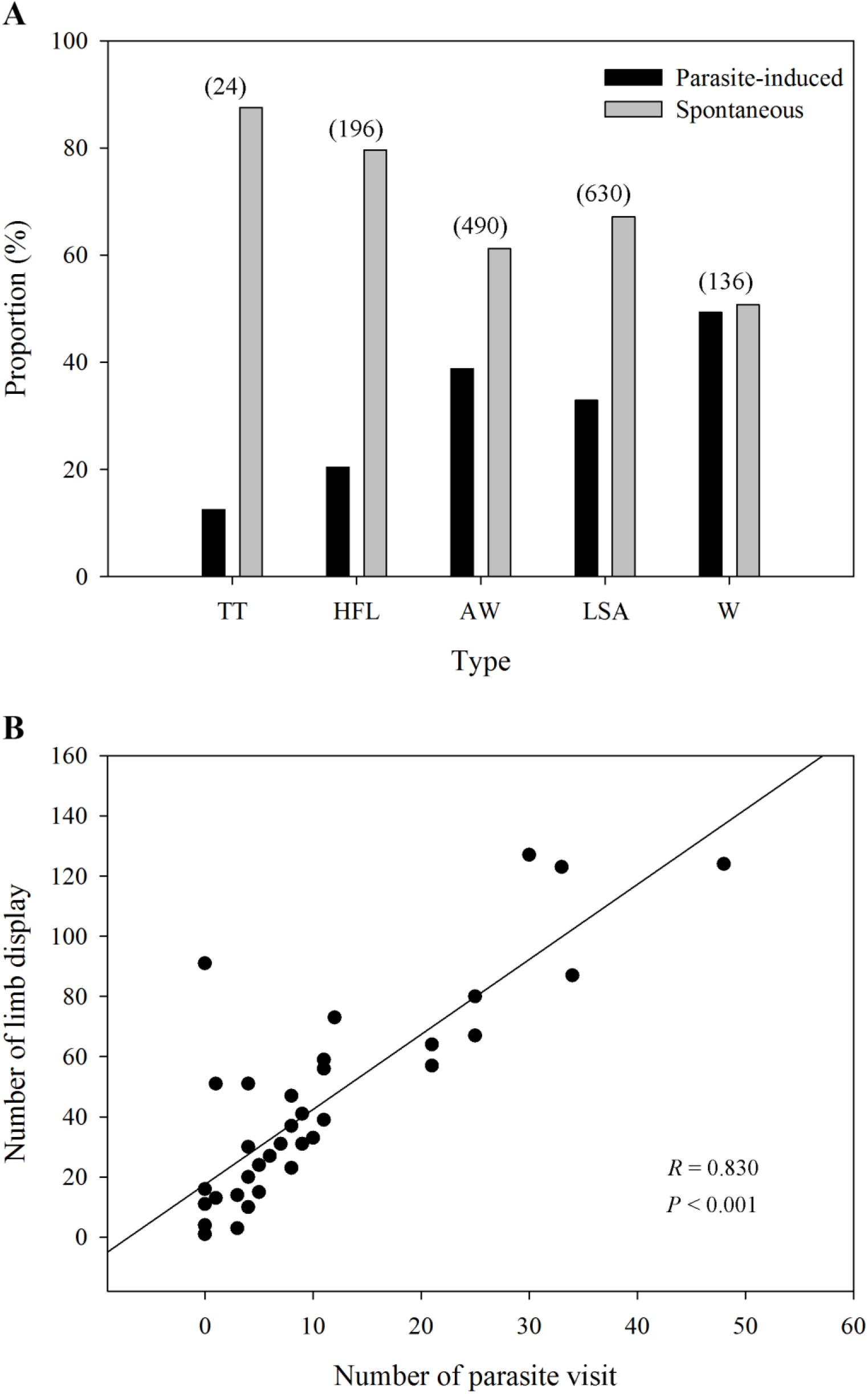
(*A*) The distribution ratio of parasite-induced and spontaneous displays in each limb movement (*N* = 69 males). TT, toe trembling; HFL, hind foot lifting; AW, arm waving; LSA, limb shaking; W, wiping. The numbers in brackets above each bar pairs represent the number of each movement, showing the richness of those visual displays. (*B*) The relationship between parasite stress and the number of all limb movements (*N* = 39 males).

### Calling males show more parasite-evoked limb movements

We compared the limb movements between calling individuals (*N* = 39) and non-calling individuals (*N* = 30) in the presence and absence of eavesdropping insects. Among the six types of limb movements, the TT and LS + FF were rarely induced by parasitic insects (table S1). We thus only compared the difference between calling males and silent males for the other four visual displays. We found calling males to produce more defensive AW (Wilcoxon rank sum test; *W* = 895; Holm-adjusted *P* < 0.001), W (*W* = 872.5; Holm-adjusted *P* < 0.001), LSA (*W* = 755; Holm-adjusted *P* = 0.032) and HFL (*W* = 736.5; Holm-adjusted *P* = 0.053) display than silent males (figure 3), presumably because calling individuals attracted more parasites than silent individuals. We also compared the whole number of parasite-evoked movements between calling males and silent males. The result suggested that calling individuals have more parasite visits than non-calling individuals (Wilcoxon rank sum test; *W* = 900.5; *P* < 0.001).

**Figure 3.**
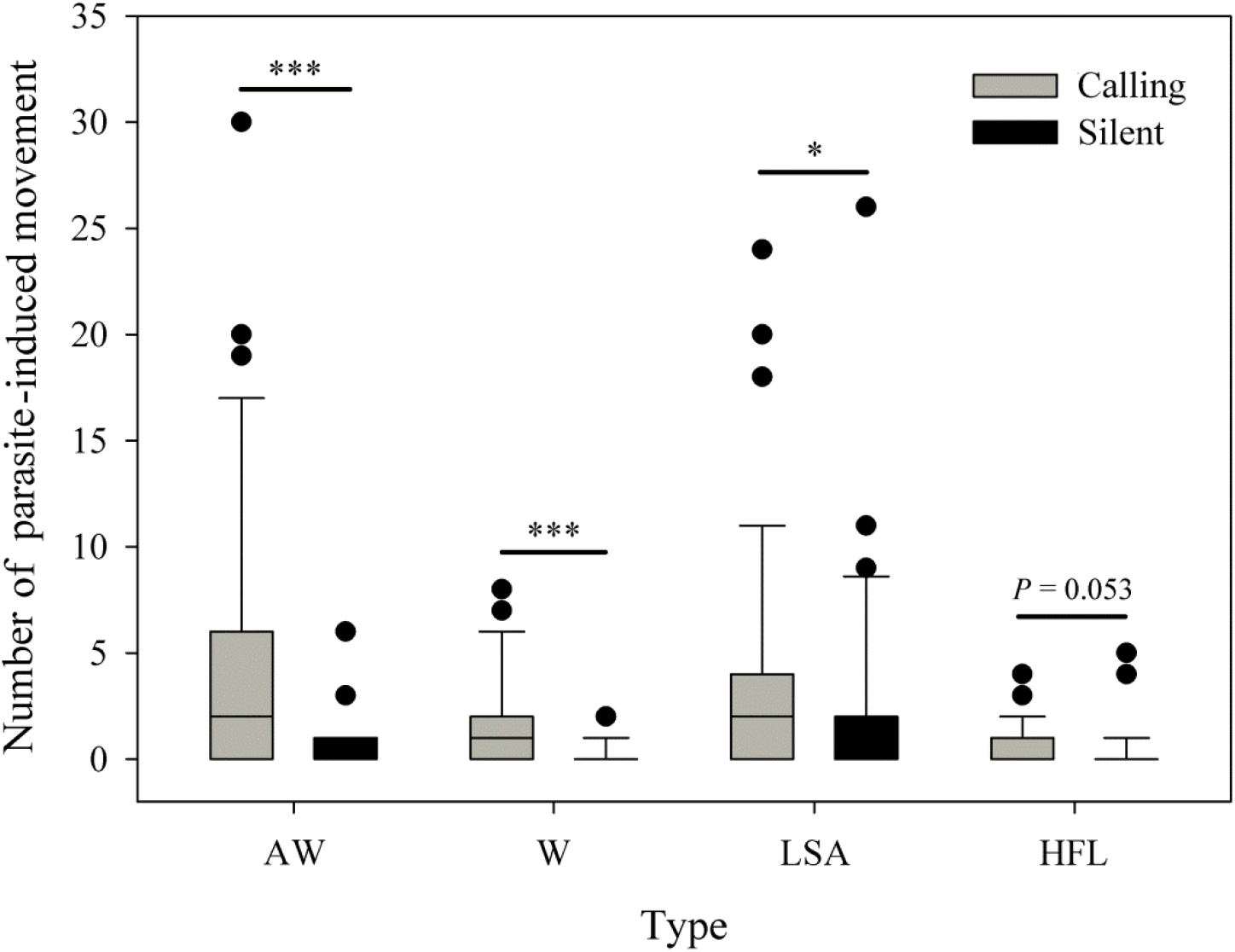
Calling males (*N* = 39) show more parasite-evoked limb movements than silent males (*N* = 30). AW, arm waving; W, wiping; LSA, limb shaking; HFL, hind foot lifting. ^*^Holm-adjusted *P* < 0.05, ^***^ Holm-adjusted *P* < 0.001.

### The role of parasite-evoked limb displays on female choice

Females expressed a strong preference during audio-visual playbacks to approach a male that performed an AW (probability = 0.81; *N* = 16; *P* = 0.021; figure 4*A*) or HFL (probability = 0.76; *N* = 21; *P* = 0.027; figure 4*B*) movement when compared to a motionless male (static control). Females did not express a preference for the W display (probability = 0.48; *N* = 31; *P* = 1; figure 4*C*) and the LSA display (probability = 0.57; *N* = 28; *P* = 0.572; figure 4*D*) during audio-visual stimulus presentation. Furthermore, females did not express a preference for the AW/HFL stimuli (dynamic visual stimuli) versus motionless stimuli (static visual stimuli) in the absence of an advertisement call (probability = 0.47; *N* = 19; *P* = 1). The LS + FF were spontaneous displays (not induced by parasites), and therefore not tested in this experiment.

**Figure 4.**
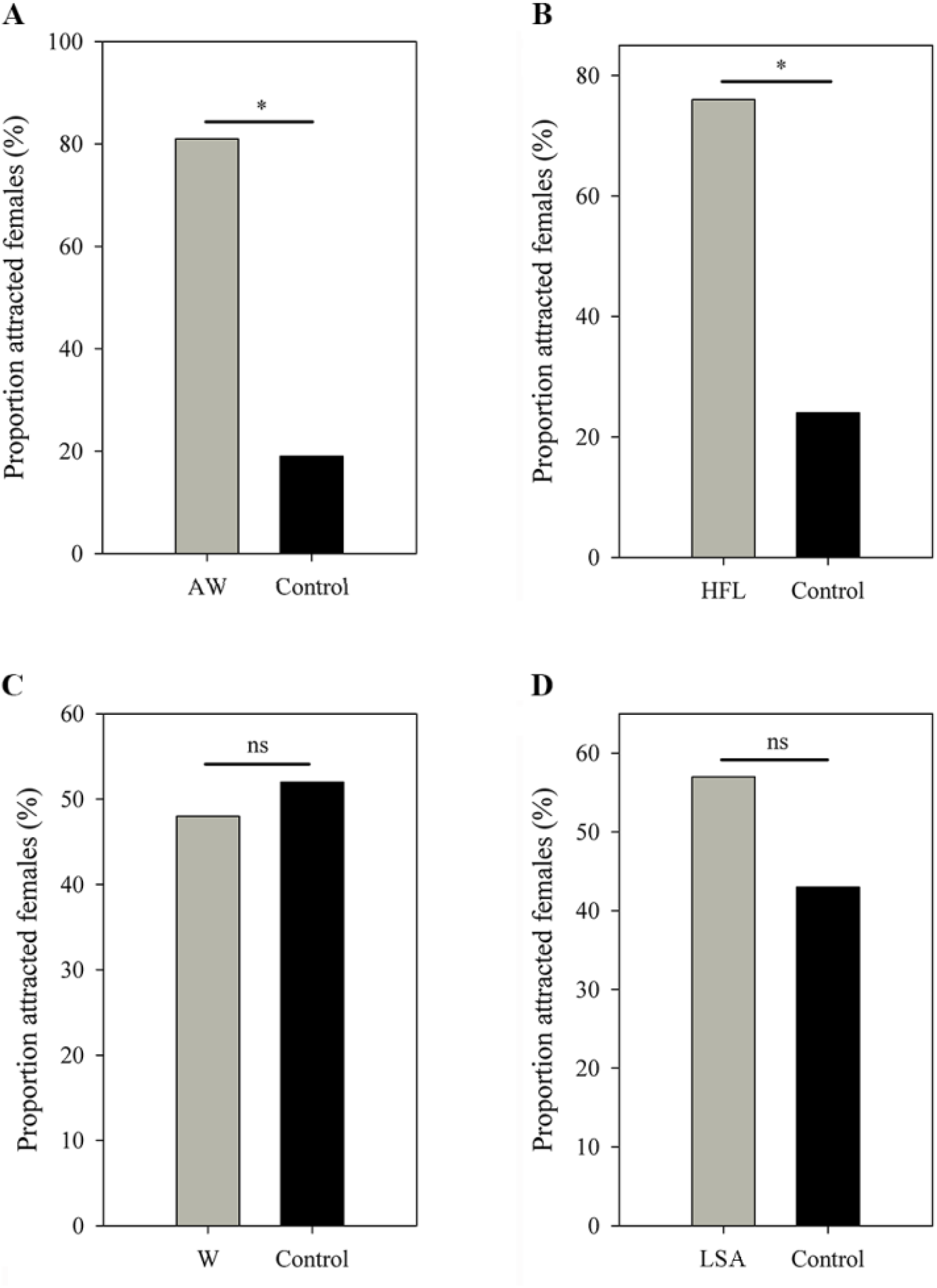
Female choices in (*A*) AW versus Control, (*B*) HFL versus Control, (*C*) W versus Control and (*D*) LSA versus Control. All limb display videos are accompanied by advertisement call and male movement, while the controls contain the same call and frog but in absence of movement. AW, arm waving; HFL, hind foot lifting; W, wiping; LSA, limb shaking. ^*^*P* < 0.05. ns, not statistically significant.

### Males show more exaggerated displays when females appear nearby

A complete recording of male-female interactions was quite difficult in the wild, because this behavior was rarely observed and frogs frequently moved among stones. Over the past two breeding seasons, we were only able to obtain four recordings of male-female interactions. For those recordings, we only analysed the LS and FF displays. These data suggest that males use more exaggerated displays (i.e. LS + FF) when females are nearby. We found a higher proportion of males to produce FF and LS displays (the most exaggerated limb movements in little torrent frogs) when females were nearby (Fisher’s exact test; *N*_*1*_ = 4, *N*_*2*_ = 39; *P* = 0.001; figure 5*A*). Only five out of thirty-nine recorded males emitted LS + FF movements when females or other males were not around (i.e. long range signaling), while all recorded individuals had such movements when females appeared nearby (i.e. close range signaling). Moreover, males produced more of those exaggerated displays when females were in close range compared to long range distance (Wilcoxon rank sum test; *N*_*1*_ = 4, *N*_*2*_ = 39; *P* < 0.001; figure 5*B*).

**Figure 5.**
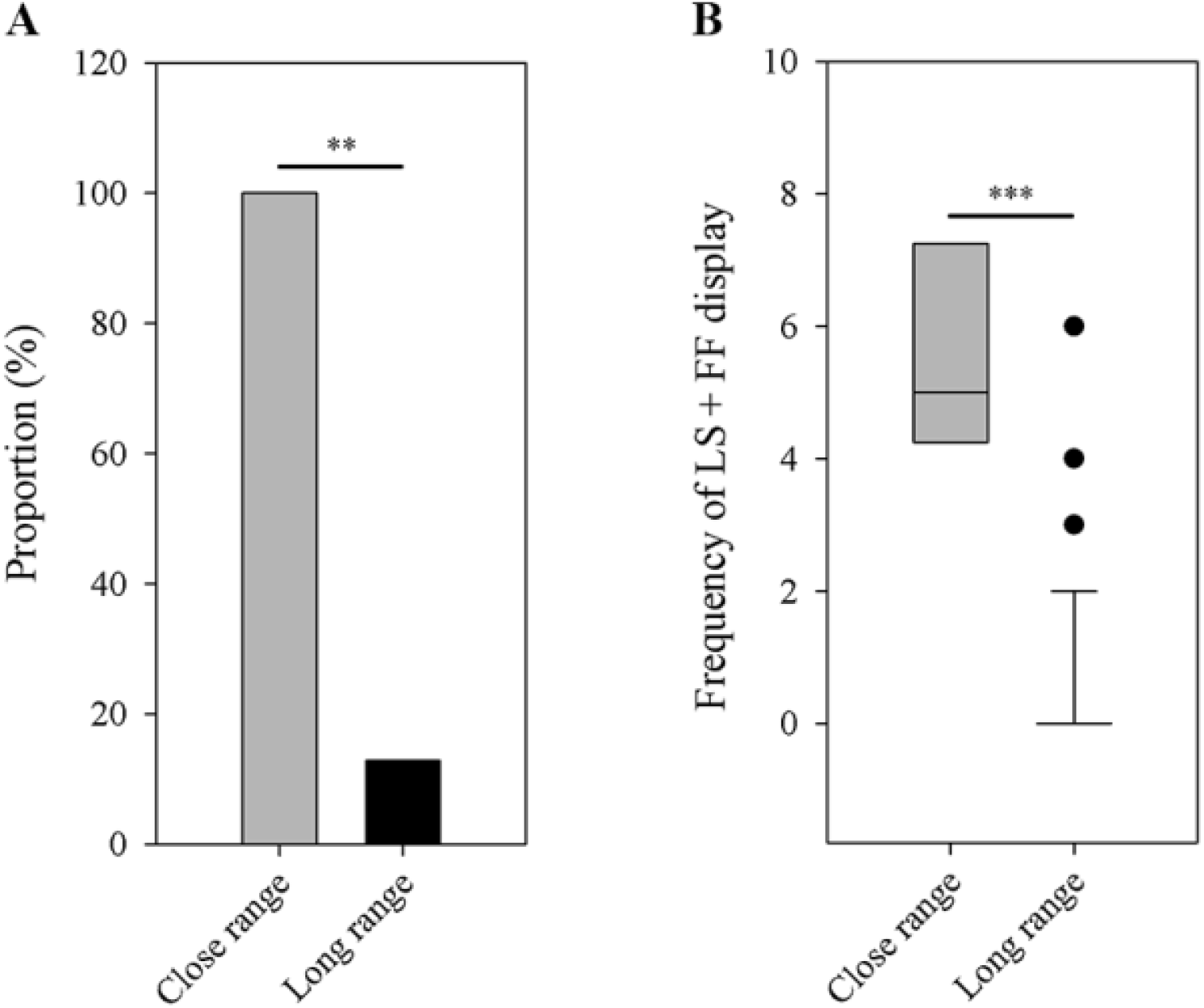
Differences between close-range group (*N* = 4) and long-range group (*N* = 39) in the number of male frogs that have the LS + FF movement (*A*) and the frequency of LS + FF display (*B*). LS, leg stretching; FF, foot flagging. ^**^*P* < 0.01, ^***^*P* < 0.001.

## Discussion

Male little torrent frogs show a rich repertoire of limb displays. By performing female choice tests and observing male-female interactions in the field, we show that several of these limb displays (AW, HFL, LS + FF) can serve as visual signal component and influence female mate-choice. All limb movements could be spontaneously generated, but their rates were increased by call-induced parasite presence. We also found calling males to produce more limb displays than silent males. Such parasite-host interactions may have important evolutionary consequences because females in our choice tests showed a strong preference for males that emitted advertisement calls accompanied by parasite-evoked movements (e.g. arm waving and hind-foot lifting). We did not find such preference for limb movements in the absence of call playback, demonstrating that these visual components are part of a multimodal display.

More and more studies show that streamside-breeding anurans often have complex physical displays including limb display as well as body display. For example, a recent investigation found that male Brazilian torrent frogs (*Hylodes japi*) have eighteen distinct physical displays which may be associated with different social contexts ^27^. Observations of Bornean ranid frogs (*Staurois guttatus*) suggest that males show abundant physical displays which are daytime- or nighttime-dependent ^29^. In little torrent frogs, we did not observe a potential visual signal related with body displays. This species, however, performed diverse common limb displays.

Stream-breeding frogs often possess abundant physical movements, but their communicative functions have rarely been tested ^14^. Most previous work analyzed those physical movements based on field observation or video, and few researchers have experimentally explored the effect of these visual displays on female mate choice. In the present study, we examined whether limb displays can serve as sexual signal and influence female choice in little torrent frogs. By video playbacks, we found females to show a significant preference for males displaying with arm waving and hind-foot lifting as compared to motionless males. Foot-flagging displays are widespread in anurans, and their functions are mainly explored from the perspective of male-male competition. For instance, male *S. guttatus* significantly increases such display when conspecific advertisement calls are broadcasted from 1-2 m distance in the field ^29^. In our study, males generated more conspicuous leg stretching and foot flagging displays when females were at close range in the field. Together, our results show that some limb movements allow male little torrent frogs to increase their attractiveness of acoustic signals to females. Color, vocal sac and physical movement, as visual subcomponents, can be added to acoustic signals in anuran multimodal communication ^30^. For example, agonistic behaviors of male *Epipedobates femoralis* are only evoked when males are exposed to vocal sac pulsations combined with acoustic signals during playback experiments ^31^. Female little torrent frogs have been shown to prefer the vocal sac movements synchronized with calls over calls only ^32^. In this study, we not only demonstrated that parasite-evoked physical movements (i.e. AW and HFL) can increase the attractiveness of acoustic signals, but also showed that males significantly increased LS + FF movement when females appeared nearby in the wild. Furthermore, males also increased such movements in response to male-male interactions (L. Zhao et al., unpublished data), which is similar with results in some other species such as *Staurois parvus* ^33^. So, both the vocal sac and conspicuous movements (i.e. AW, HFL, and LS + FF) are visual subcomponents of multimodal displays in little torrent frogs.

Frogs and toads are known to be able to see well in dim-light ^34^. Moreover, many anurans perform behaviors only at environmental illuminations with very low levels ^35^. Due to their phototactic responses ^36^, some video playback studies may have obtained odd results in the past. Little torrent frogs are diurnal species who can communicate with acoustic and visual signals under high-light as well as darkness environments. In order to avoid possible effects from light, we set a low environmental illumination according to a natural level. In addition, we argue that little torrent frogs actually recognize the video image as a conspecific stimulus. There are two reasons for this. First, females discriminated between moving and stationary stimuli. Second, more conspicuous movements (e.g. HFL) had greater attractiveness than less conspicuous movements (e.g. limb-shaking).

Among four visual movements in our playback experiments, the attractive AW and HFL are more conspicuous than the unattractive W and limb-shaking because the display of AW and HFL involves more occupation in terms of time and space. More conspicuous physical movements increase receiver’s attention ^37, 38^, so the difference in conspicuousness may be related to the attractiveness of these movements. Such speculation is further supported by our analysis on LS and FF displays. The two movements are the most conspicuous visual signal in little torrent frogs. Males significantly increase the two movements when females appeared nearby in the wild. Such conspicuous displays, however, are not used during parasite interactions because male frogs generally perform defensive motions when parasites fall on their body (or limbs) or fly very close to their body (or limbs). Under these conditions, small and medium limb movements are sufficient, while the exaggerated motions (LS and FF) seem to be unnecessary. Therefore, the LS and FF displays are specially performed for conspecific communication, but may have evolved out of the less conspicuous parasite-induced limb movements.

The evolution of dynamic visual signals is often influenced by several social and environmental factors such as territorial aggression ^16, 39, 40^, diurnality ^13^, background noise ^41, 42^ as well as predation pressure ^43^. Visual movements or movement-involved multimodal signals can result in senders with more chance of being seen by predators or parasites. Physical displays thus often need to increase the attention of intended receivers while limiting the eavesdropping of unintended receivers ^44^. In little torrent frogs, male limb displays increased with parasite stress, and such defensive movements can serve as visual cues. These results suggest that parasite stress can induce more visual movements and increase the complexity of audiovisual multimodal displays in little torrent frogs as a by-product. In this study, calling males had a larger parasitic risk than silent males, and we identified *Corethrella* spp midges which are known to localize frog hosts by eavesdropping on their calls ^24^. However, more studies are needed to further examine whether and how the species is eavesdropped by sound-locating midges.

The idea that some physical movements are the raw material of dynamic visual signals has been proposed for many years ^13, 15^. In the Panamanian golden frog (*Atelopus zeteki*), for instance, the conspicuous semaphore signal is supposed to originate from a standard stepping motion ^45^. However, it is largely unknown how physical movements are incorporated into communicative systems. Sexual selection is believed to be an important driver and the evolution of physical movements may be favored when females are sensitive to some movements in specific environments ^46^. During courtship, complex signals are often preferred by females and males frequently include sexual displays more than one channel ^1, 47^. So physical movements from males may be incorporated into multimodal communication systems if they tap into the sensory bias of females and are beneficial to increase the attractiveness of males (sensory exploitation process ^48^). In our study, females little torrent frogs showed a significant preference for the conspicuous defensive movements when the advertisement calls were simultaneously broadcasted. In noisy streams, acoustic cues plus the relative conspicuous movements may benefit animals to overcome auditory masking by flowing water ^49^. Our study experimentally shows that such incorporation of non-sexual movements may actually work to increase female preference and thus becomes part of a multimodal display.

The simplest movements (LS/W) are hardly used during parasite interactions and do not induce female preference. The intermediate movements (AW/HFL) are used during parasite interactions and induce a preference. The most complex movements (LS+FF) are only used during male-female (or male-male) interactions. This observed pattern could therefore reflect the evolutionary history of the visual display, from a simple to an advanced stage, where the most complex movements are no longer used in their original context (parasite defense) but only for their new function (sexual communication). Interestingly, in many anurans such as e.g. *Micrixalus saxicola* and *S. parvus* ^33^, the most complex display also involve foot-flagging.

In conclusion, we show that calling behaviors and the levels of parasite interference are correlated and males produce diverse defensive limb movements in order to avoid those unintended receivers in little torrent frogs. By female mate-choice tests, we find the relative conspicuous defensive movements increase the attractiveness of male calls to female frogs. Thus, we suggest that parasite-host interaction may increase the complexity of audiovisual sexual displays. For the first time, we experimentally demonstrate that movements evoked by interspecific activities may evolve via increasing female preference. Our results, together with phylogenetic studies in future, would increase our understanding towards the evolutionary origin of dynamic visual cues and the relationship between natural or sexual selection pressure and multimodal communication behavior.

## Methods

### Field site and study species

The study was carried out in Wuzhishan Nature Reserve (18°55’N and 109°41’E), Hainan Province, China. Average annual rainfall and air temperature in this area are 1800-2000 mm and 22.4°C, respectively. We focused on the little torrent frog, a species that lives in mountain streams of tropical forests accompanied with high-level torrent noise. Little torrent frogs produce a simple advertisement call (∼5.5 s) consisting of a series of repeated notes (∼50 notes/call) in which each note (∼45 ms) has the dominant frequency around 4 kHz ^50^. Male visual displays, however, are very complex and involve vocal-sac inflation, various limb movements as well as other physical movements. Vocal sac inflation always accompanies call production (obligatory coupled signal components), whereas limb movements can be made simultaneous as well as independent from calls (flexibly coupled constituent parts). Likewise, a previous study has showed that the vocal sac inflation is a part of multimodal signals ^32^.

Field data (visual displays and ecological factors) was obtained in May-July 2017 and August-September 2018, and female choice tests were conducted in July-August 2019 in our lab in Wuzhishan. Little torrent frogs can breed and communicate with visual (limb movements) and acoustic signals (calls) day and night. Male videos were recorded at day in order to obtain better frames, while females were collected and tested at night in order to assure the environmental conditions (e.g. light condition) between behavioral room and field were similar during the experiment. Gravid females (characterized by a plump abdomen) were collected in a stream around the management station of Wuzhishan Nature Reserve, between 8PM and 11PM. These females were brought to the lab in containers which included some water and rocks from their capture sites. Prior to the test, all individuals were placed in a quiet and dark environment for at least 1 h. In order to avoid repeated testing, individuals were toe-clipped and released at their capture site on the same night after the test. All procedures were approved by the management office of the Wuzhishan Nature Reserve and the Animal Care and Use Committee of the Chengdu Institute of Biology, CAS (CIB2017050004 & CIB2019060012).

### Behavioral recordings and analyses

We searched for focal males in a stretch of stream (∼1.5 km), starting at the station of the Wuzhishan Nature Reserve and ending at the source of the Changhua River. In order to control for differences in temperature, light and daily rhythm, we only recorded calling males versus silent males, between 10AM and 12PM (i.e. 2 hours), on sunny days. This species is sexually dimorphic, and females have larger body size and width-length ratio than males. We identified them according to those morphological characteristics. Thirty-nine calling frogs and thirty silent frogs were continuously recorded for 10 min using a video camera (GZ-MG465BAC, JVC, Kanagawa, Japan) from a distance of 0.3-0.5 m. Their calling behaviors and visual displays are not apparently disturbed by our recordings at such distance, because they always stayed at the original location and performed audio or visual signals as the period prior to be recorded. After each test, the temperature and humidity were measured with an electronic thermohygrometer (YHZ-90450, Yuhuaze, Shenzhen, China) and the background noise (Z-weighted) was measured near the frog’s head via a sound level meter (AWA 6291, Hangzhou Aihua Instruments, Hangzhou, China) pointing in vent-snout orientation. We compared the noise levels between calling individuals and silent individuals. The data revealed that the background noise did not differ significantly in two groups (Wilcoxon rank sum test; *N*_*1*_ = 39, *N*_*2*_ = 30; *W* = 567.5; *P* = 0.837).

We also filmed males that successfully attracted females to close-range and interacted for about 10 min. Males with a female close by (less than 0.5 m) and those without one (without female or other male within 1.5 m) were defined as the close-range and long-range categories, respectively. The limb displays were classified into seven types according to two published ethograms for anurans (see Results). When midges or mosquitoes were on the body of frogs, they would shake their body or move their limbs to repel those parasites (movie S1). Frogs also produced limb movements towards parasites that were flying close to their body (movie S2). We determined whether or not a limb movement was directly evoked by an insect according to above foraging behaviours. For all frogs, we watched the whole video and scored the number of each spontaneous movement as well as the number of each display that was evoked by midges or other insects (i.e. insect-induced movement). The number of passive motions was used to represent the level of parasite interference. Among seven movements, the LS and FF were not produced during parasite interactions. For the two conspicuous movements, we only included the number of spontaneous type.

### Mate choice tests

Video stimuli used in our mate choice trials were filmed with a camera (D7100, Nikon, Tokyo, Japan), mounted to a tripod, connected to a directional microphone (MKE400, Sennheiser, Hanover, Germany). All stimuli were recorded in front of focal frogs (*N* = 20 males) from a distance of 0.3-0.5 m. We identified four types of limb movement that were often evoked by interactions with parasites. For each type (arm-waving, foot-lifting, wiping or limb-shaking), we selected three representative videos, from three different frogs, containing a sequence in which a limb movement was accompanied by a nearby flying midge, and a sequence in which only a flying midge occurred (but with no limb movement). Both sequences were in absence of vocal sac movement. There were two reasons for the exclusion of vocal sac. First, the movement of vocal sac is flexibly coupled with limb displays in natural conditions. Second, the role of limb movements may be masked by vocal sac movement because it has a strong sexual attractiveness and can play a role in mate choice when coupled with advertisement calls ^32^. In addition, the LS and FF displays were not produced during parasite interactions. We therefore did not include them when examining the roles of parasite-evoked movements in female mate-choice.

The video stimuli were edited in Adobe Premiere Pro CS6. We firstly cut each clip to 6 s and replicated them to generate a new video with a total length of 10 min, respectively. In those videos, the display rate of each movement was within a natural range. Next, we changed the audio channel of the video by replacing the original recording with standard sound files. The standard sound files (with flowing water and calls included) were produced according to an stimulus used in a previous study ^51^. The stimulus was synthesized based on the characteristics of thirteen males (average dominant frequency: 4.3 kHz) and three typical places of flowing water. In little torrent frogs, the call rate of advertisement calls varies from 0.61 calls/minute to 3.03 calls/minute ^50^. In this study, call rate was set to a low level (1 call/minute) in order to best simulate conditions in which visual displays might be exploited. For each kind of display, we constructed three audio-visual stimulus pairs (from three different males) always containing a video with a limb display and a video without display for each type of limb movement. In those stimuli, calls were partly overlapped with limb displays on the videos, which were similar with natural scenarios. Moreover, we re-used three videos to produce three stimulus pairs without the advertisement calls (type 5; details are included in Results). Taken together, five types of audio-visual stimuli with a total of fifteen pairs were constructed in this experiment.

We tested female preferences for our stimulus pairs in a sound-attenuating phonotaxis chamber (1.5 × 1.5 × 1.2 m, L × W × H) under infrared lighting. We placed a LCD monitor (17S4LSB, Philips, Amsterdam, Netherlands) and a speaker (JBLCLIP + BLK, JBL, Los Angeles, USA) at each side of two nearby corners, which were used to present sounds and frames, respectively. Such method with frogs in videos has successfully been used to test for sexual preferences in previous studies ^52, 53^. Two monitors were moved and rotated to ensure that both the distance between the two monitors and the distance between the initial female placement point were all 1 m, resulting in a 60° angle between two frames with respect to the initial female placement position (figure 1). All males were life-sized in the screens. The brightness and color of screens were calibrated using light meter (TES-1399, TES, Taibei, China) and color correction instrument (Spyder5 ELITE, Datacolor, Lawrenceville, USA), respectively. Moreover, the speakers were fixed right under the frame presentation areas during playback (figure 1).

For each female, the sound pressure levels of both speakers (SPLs) were calibrated with a sound level meter such that calls were 75 dB (re 20 μPa) at the initial female placement position. Such intensity is near the auditory threshold ^51^, and was set to increase the likelihood of both calls and movements being noticed ^54, 55^. Prior to each playback, a piece of black sponge was placed in front of female frog in order to avoid a possible interference (i.e. light or other visual information prior to each playback) from the screens. The start of each playback was simultaneously conducted with the removing of the sponge during the experiment (females do not move without sound or video playback). A choice was scored when a female approached a speaker-monitor combination within 5 cm. We considered a female as lacking motivation if she failed to make a choice within 10 min. For each frog, we presented the screen with a visual movement (a moved male) versus the screen without the visual movement (the same male not moving) to examine the role of visual displays in absence of acoustic signals. We also presented the screen with audio-visual (call plus movement) versus the screen with audio (call plus same male not moving) stimulus pairs to test the role of visual displays in presence of acoustic signals. All females thus experienced both movement/no-movement and audio-visual/audio conditions. In order to avoid potential side effects, each stimulus pairs on the left versus right sided monitor were randomly broadcasted during the experiments. In order to avoid potential sequence effect, the order of different stimulus pairs was randomized across females. Besides, females could finish multiple tests sequentially and they were not given a break during playback experiments.

### Data analyses

We analyzed all data on male visual display and female mate-choice in R (v.3.5.3). We used a Pearson correlation analysis to determine the relationship between the level of parasite interference and the number of visual display. We carried out Wilcoxon rank sum tests to evaluate the difference of parasite-induced visual displays (i.e. limb-shaking, wiping, arm-waving and hind foot lifting) between calling frogs and silent frogs (quantity of each per 10 minutes). We also used the Wilcoxon rank sum test to determine the change of male exaggerated foot-flagging and leg-stretching displays when female was around. In these tests, multiple comparisons were corrected by adjusting *P*-values using Holm’s method. Moreover, we employed Fisher’s exact test to examine whether males emitted more exaggerated foot-flagging and leg-stretching displays when females appeared nearby. Female mate-choice was compared with a binomial test. *P* < 0.05 was considered statistically significant.

## Acknowledgments

We thank Yanlin Cai, Xiaoqian Sun and Xiaofei Zhai for their help during the wild recordings. This work was supported by National Natural Science Foundation of China (31772464), Youth Innovation Promotion Association CAS (2012274) and CAS “Light of West China” Program.

## Data availability

Data used to generate the results are available from the Dryad Digital Repository: https://doi.org/10.5061/dryad.f1vhhmgzg.

## Supplementary 1

**Figure S1.**
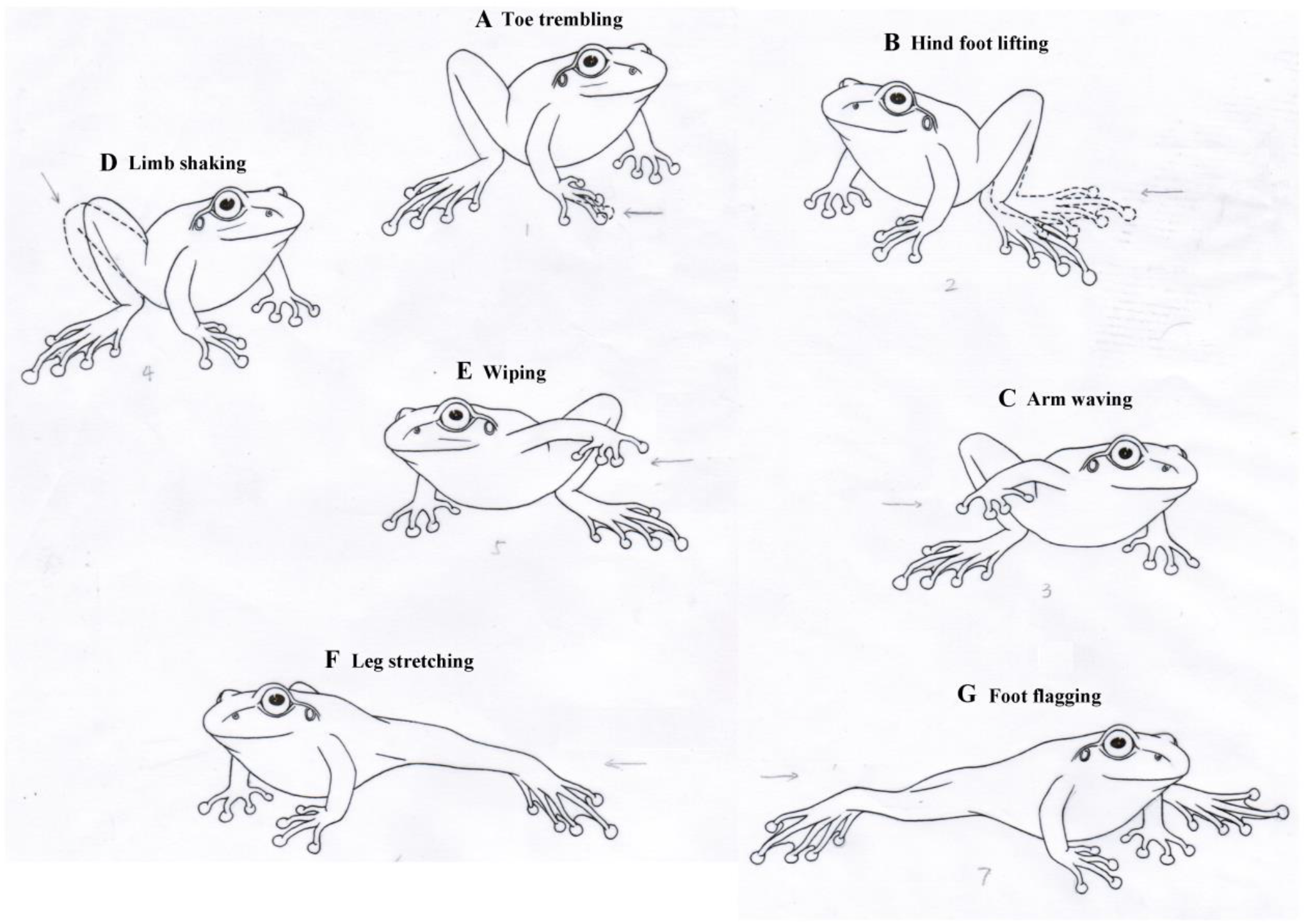
Examples of different limb motion displays.

**Figure S2.**
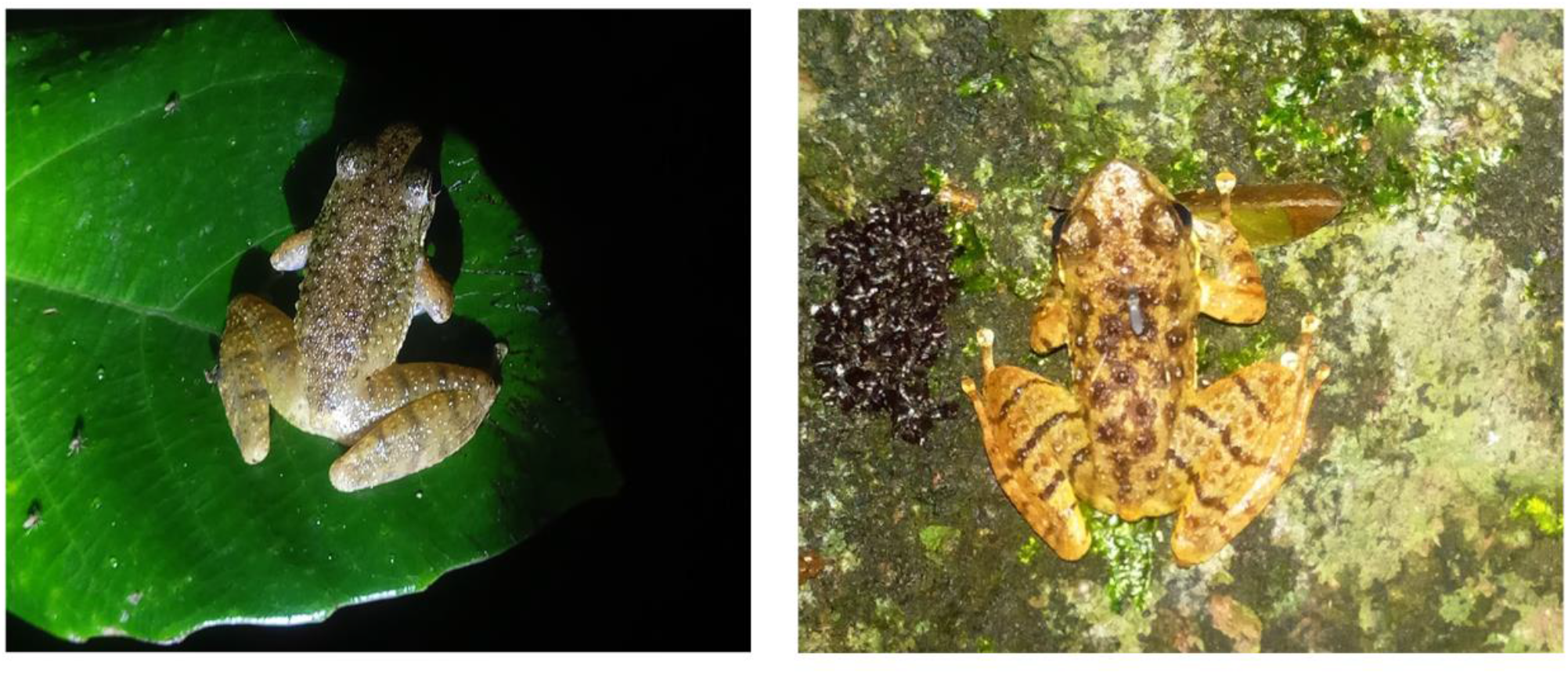
Photos that frogs are being bitten by potential parasites.

## Supplementary 2

**Movie S1**. Video that frog produces defensive motions in order to repel midges.

**Movie S2**. Video that frog produces movement towards a flying parasite.

## Supplementary 3

**Table S1**. The data of parasite-induced and spontaneous displays in each limb movement for calling males, silent males and males that have females nearby.

